# Kernel machine tests of association between brain networks and phenotypes

**DOI:** 10.1101/341495

**Authors:** Alexandria M Jensen, Jason R Tregellas, Brianne Sutton, Fuyong Xing, Debashis Ghosh

## Abstract

Applications of quantitative network analysis to functional brain connectivity have become popular in the last decade due to their ability to describe the general topological principles of brain networks. However, many issues arise when standard statistical analysis techniques are applied to functional magnetic resonance imaging (fMRI) connectivity maps. Frequently, summary measures of these maps, such as global efficiency and clustering coefficients, collapse the changing structures of graph topology from many scales to one. This can result in a loss of whole-brain spatio-temporal pattern information that may be significant in association and prediction analyses. Drawing from the electrical engineering field, the resistance perturbation distance is a quantification of similarity between graphs on the same vertex set that has been shown to identify changes in dynamic graphs, such as those from fMRI, while not being computationally expensive or result in a loss of information. This work proposes a novel kernel-based regression scheme that incorporates the resistance perturbation distance to better understand the association with biological phenotypes from fMRI using both simulated and real datasets.

## Introduction

Since its introduction in the early 1990s [1] [2], functional magnetic resonance imaging (fMRI) has rapidly grown to become the most popular technique to observe the living human brain, with nearly 30,000 papers published on the topic in 2015 alone [3]. First described by Ogawa in 1990, the blood oxygen level dependent (BOLD) signal is the most common method of fMRI [4]. The BOLD signal employs hemoglobin as an endogenous contrast agent, exploiting the magnetization difference between oxy- and deoxyhemoglobin to create the fMRI signal [5]. This BOLD signal is theorized to be comprised of three determinants: a neuronal response to either an endogenous or exogenous stimulus, the relationship between the neuronal and hemodynamic responses, and the hemodynamic response itself [5]. MRI measures this neuronal responses indirectly through its hypothesized hemodynamic correlate, in contrast to EEG and MEG, two popular techniques for analyzing functional connectivity in the brain that rely on quantification of electrical activity. In addition to the differences in focus of these techniques, resolutions differ. While EEG and MEG provide temporal resolution on the scale of milliseconds at the cost of spatial localization ambiguity; fMRI has much higher spatial resolution but is limited in temporal resolution due to the slower rate of brain hemodynamics [6]. While all of these techniques offer noninvasive ways to study brain activity, fMRI has quickly outpaced EEG and MEG in popularity within the research community.

Techniques like fMRI can be used in biomedical research to examine localization of brain regions engaged by a particular task, determining brain networks, and predicting psychological or disease states [6]. While most fMRI studies initially focused on the examination of brain regions engaged during a specific task, increased attention has been paid examining the connectivity of the entire brain at rest, commonly referred to as resting state fMRI (rs-fMRI) [7]. Analysis of rs-fMRI can help yield information about the strength of connections within and among brain regions that may be unique to clinical populations [7] [8]. All of these objectives can be achieved through the application of statistical techniques that address the specific complications that arise from analysis of spatially correlated, four-dimensional data.

The most common technique used to analyze fMRI data is a mass univariate analysis (MUA). In MUA, a general linear model is fit at each voxel independently with a combination of experimental conditions and biological confounders as covariates [9]. This creates a map of parameter estimates and test statistics that is then thresholded to identify significant voxels; the location and clustering of these significant voxels inform the functional relationships within the brain. Time series methods have also been successfully implemented to examine the interregional correlation values over the length of the fMRI scan [7]. However, these techniques ignore the underlying spatial relationships within the data; even though voxel responses are correlated, mass univariate analysis and its time series extension do not fully account for the underlying spatial correlation [9]. However, by extending the general linear model framework to allow for the modeling of non-linear relationships, more complex associations can be fit. One such way to accomplish this is through the use of kernels, which are weighting functions used to estimate the conditional expectation of a random variable [10].

In contrast to the use of general linear models and kernel regression, applications of quantitative network analysis through graph theory have become popular in the last decade due to their utility in describing the general topological principles of brain networks [11]. First introduced by Leonhard Euler in his 1736 solution to *Seven Bridges of Konigserg,* graph theory can be used to model a range of relations and processes in physical, biological, social, and information systems [?]. The application of graph theory to study the underlying structural and functional connections within the brain was first introduced by Ed Bullmore and Olaf Sporns in their seminal work, published in 2009 [11]. In their work, Bullmore and Sporns showed how connectivity analyses can be used not only to analyze structural networks that represent the architectural connections within and between regions, but also to analyze the underlying functional networks that can elucidate how this architecture supports various neurophysiological dynamics [11]. Numerous studies have reported that brain network parameters, derived from fMRI, EEG/MEG, and structural MRI, differ between subjects based on task or underlying biological or physiological condition [11]. However, many issues arise when applying standard statistical methods to fMRI connectivity maps. Frequently, summary measures of these maps, such as global efficiency and clustering coefficients, collapse the changing structure of graph topology from many scales to one [12]. This can result in a loss of whole brain spatiotemporal pattern information that may be significant in association and prediction analyses.

This study proposes a kernel regression scheme that incorporates the resistance perturbation distance to better predict biological phenotypes from fMRI using both simulated and real datasets. Drawing from the electrical engineering field, the resistance perturbation distance (RPD) is a quantification of similarity between graphs on the same vertex set that has been shown to identify changes in dynamic graphs, such as those from fMRI, while not being computationally expensive or result in a loss of information [12]. By incorporating the RPD into a kernel distance function, the high-dimensional feature space of brain networks, defined on input pairs, can be generalized to non-linear spaces; this allows for a wider class of distance-based algorithms, rather than the restrictive squared distance, to be representative of the similarity between two networks. We hypothesize that this algorithm will show significant associations between the metric and phenotype.

The remainder of this paper is organized as follows. The methods section will describe the derivation and properties of the resistance perturbation distance, the general framework for distance-based kernels, and a kernel-based score test. We then apply these methods under a variety of simulation paradigms and to the COBRE-I dataset. Finally, we conclude the paper with a discussion and proposal of future directions. All R code will be made available as supplementary material to this manuscript.

## Materials and methods

### The resistance perturbation distance

Like many complex systems, the human brain is highly dynamic, where the relationship between regions changes with respect to time. The brain is a highly plastic organ, able to reorganize itself through modifications of its neuronal connections. This feature is unique to the central nervous system: neuroplasticity occurs at the beginning of life, where the immature brain organizes itself, when it is subjected to trauma or injury, and throughout adulthood whenever something new is learned. Neuroplasticity also relates to the brain’s ability to have both specialized and integrated regions, known as neuronal clusters. Depending on the needs of the brain, neuronal clusters, especially those deemed to be highly influential [13], communicate across many regions of the brain. The most popular graph theory measures collapse this complex system from many scales to one, resulting in a loss of information. In contrast, the resistance perturbation distance is a quantification that is flexible enough to account for configurational changes that can occur on a local scale - through local neighbors on the same node - or on a global scale - through connections between clusters or hubs [12].

Let *G* = (*V, E, w*), be an undirected, weighted graph that is connected (there is a path between every pair of nodes) and contains no self-loops (edges that connect nodes to themselves), where *V* = {1, 2,…, *n*} is the vertex set, *E* is the edge set, and *w* is a symmetric weight function that provides a quantification of the strength of the relationship between two vertices. The higher the value of *w,* the stronger the relationship between two vertices *i* and *j.* If we define the weighted adjacency matrix to be

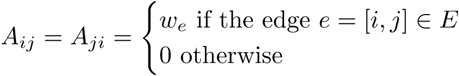

the degree matrix’s definition is simply 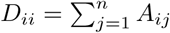 [12]. Using the adjacency and degree matrices, the Laplacian matrix, **L,** is a symmetric and positive semi-definite matrix, where **L** = **D** − **A** [12]. The Lapacian matrix has many special properties, including that it can be spectrally decomposed into eigenvectors and eigenvalues; as **L** is positive semi-definite, then all eigenvalues *λ_j_* ≥ 0. Further, if we find the Moore-Penrose pseudo-inverse of **L,** denoted **L**^†^, by definition of **L, L**^†^ is also symmetric.

An important aspect to these adjacency matrices is that a family of distances on *G*^(1)^ and *G*^(2)^ can be induced through the application of a matrix-to-matrix function *ϕ* on the corresponding adjacency matrices. Monnig and Meyer define a general graph distance as,

“Given a matrix-to-matrix function, *φ,* on a square matrix, *M_n×n_,*

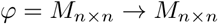

and a distance *d* on *M_n×n,_* we define the pseudo-distance *d_φ_* between two graphs *G*^(1)^ and *G*^(2)^ as

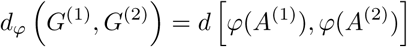

where *A*^(1)^ and *A*^(2)^ are the adjacency matrices of *G*^(1)^ and *G*^(2)^, respectively. If *φ* is injective, then *d_φ_* defines a distance [12].”

This definition is important in that it decouples two important aspects to the distance *d_ϕ_.* The matrix-to-matrix function *ϕ* extracts geometric or configurational properties from each graph while the distance *d* can be used to emphasize the relative size of variations in *ϕ.* Koutra et al. expand on this definition, defining axioms that any distance measure should satisfy:

1. *d_φ_* (*G*^(1)^, *G*^(1)^) = 0
2. *d_φ_* (*G*^(1)^, *G*^(2)^) = *d_φ_* (*G*^(2)^, *G*^(1)^)
3. *d_φ_* (*G*^(1)^, *G*^(2)^) → 0 as the number of nodes *v* → ∞, where the edge sets between *G*^(1)^ and *G*^(2)^ are complementary [14].

As well, a distance measure should satisfy the following properties:

1. Edge Importance: a change in an edge that creates disconnected components within the graph should be penalized more than changes that maintain its connectivity properties.
2. Weight Awareness: the larger the weight of a removed edge, the greater its impact on the distance.
3. Edge- “Submodularity”: a change is more meaningful in a sparse graph than in a denser graph that are both defined on the same vertex set.
4. Focus Awareness: random changes in a graph result in a smaller impact than targeted changes [12] [14].

The concept of effective resistance, commonly seen in the electrical engineering field, can be understood in terms of a commute time, which can be analogously extended to the graph theoretic measure of path length but with a richer choice of distance *d.* Monnig and Meyer showed that the effective resistance between two vertices falls under the definition of a general graph distance, but also preserves the three axioms and four properties detailed above. Because the BOLD signal measures the indirect correlate of neuronal responses in the brain, which are the transfer of action potential between neurons, seeing the brain as a circuit board is not an uncommon analogy. Therefore, the use of effective resistance can be easily extended to the summarization of fMRI data. Effective resistance can be defined as

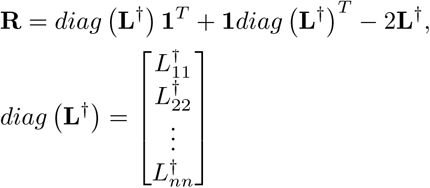

[12].

Using this, then define the resistance perturbation distance as the element-wise p-norm of the difference between effective resistances such that

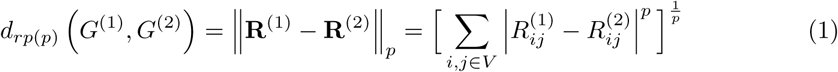

for 1 ≤ *p <* ∞ and

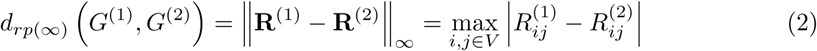

for *p* = ∞ [12].”

This distance metric defines a distance on the space of connected, undirected, weighted graphs on the same vertex set, where the effective resistance **R** is fully and uniquely defined by a graph’s Laplacian matrix **L** and, for 1 ≤ *p <* ∞, the element-wise p-norm, ||·||*_p_*, is a norm for a matrix *M_n×n._* Thus, the distance can easily be shown to be non-negative, symmetric by application of the definition of an element-wise p-norm and satisfies the triangle inequality,

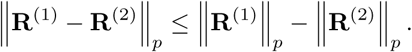

Additionally, if we observe *G*^(1)^ = *G*^(2)^, then *d_rp_*_(*p*)_ (*G*^(1)^, *G*^(2)^) = 0 because **R**^(1)^ = **R**^(2)^ by definition. Conversely, if *G*^(1)^ and *G*^(2)^ are two graphs with the same effective resistance matrix, **R**^(1)^ = **R**^(2)^, this implies the equality of the Laplacian matrices, **L**^(1)^ = **L**^(2)^ and, continuing on this train of thought, equality of their weighted adjacency matrices, **A^(^**^1)^ = **A**^(2)^ [12].

#### Distance-Based Kernels and a Kernel-Based Score Test

Rather than assume a parametric form in the relationship between functional connectivity matrices and phenotypic classification, kernel distance estimation was implemented as a non-parametric way to quantify the similarity between data instances. A range of kernel functions are used in statistics, where the choice of kernel determines the function space used to approximate the relationship between two variables [15]. The most popular is the Gaussian kernel, 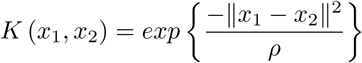, where ||*x*_1_ − *x*_2_||^2^ is defined as the squared Euclidean distances and *ρ* is an unknown bandwidth or scaling parameter. More generally, a distance-based kernel, of which the Gaussian kernel is a member, is denoted as

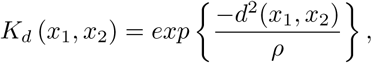

where *d*^2^ (*x*_1_*, x*_2_) is a distance function [10]. *K_d_* (*x*_1_, *x*_2_) can be thought of as a measure of similarity between two subjects *x*_1_*, x*_2_ in terms of some multidimensional variable set *Z.* This similarity measure can then be incorporated into a regression framework to test to what extent variation in *Z* can explain variation in the phenotypic outcome, *Y.*

In the case of a dichotomous outcome, assume a logistic regression framework of the semiparametric form

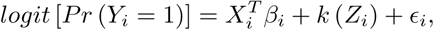

where *X_i_* is a matrix of covariates whose association to the dichotomous phenotypic outcome, *Y_i_,* is to be parametrically estimated, *k* (·) is a centered, smooth kernel function, and *Z_i_* is a vectorized form of the RPD matrix from the previous section [15] [10]. An important feature of *k* (*Z_i_*) is that it lies within a Reproducing Hilbert Kernel Space (RHKS). A RHKS is a function space if *f* and *g* are close in norm, ||*f* − *g*|| *< ϵ,* then *f* and *g* are also pointwise close, ||*f*(*x*) − *g*(*x*)|| *< ϵ_x_* for all *x.* A hypothesis test can be conducted to determine whether the multidimensional variable set *Z_i_* is associated with *Y,* controlling for *X,* of the form

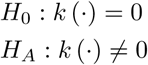

[15] [10].

Assuming that *k* (·) lies within a RHKS, 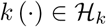, *β* and *k* (·) can be simultaneously estimated by maximizing the penalized log likelihood function

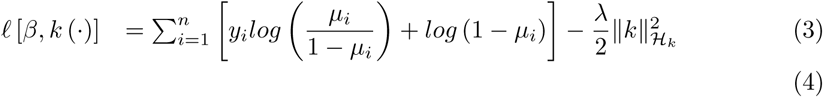

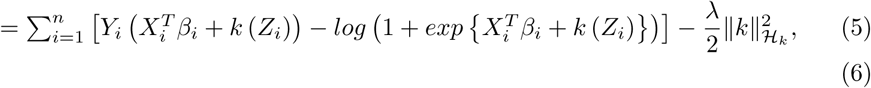

where *λ* is a regularization parameter that reflects the trade off between model complexity and goodness of fit [16]. At its boundaries, *λ* = 0 reflects a saturated model, while *λ* = ∞ reduces the model to a fully parametric logistic regression model. However, it should be noted that there are two unknown parameters within *ℓ*[*β, k* (·)]: the regularization parameter *λ* and bandwidth parameter *ρ.* Intuitively, *λ* controls the magnitude of the unknown function *k* (·) while *ρ* controls the smoothness of *k* (·) [15]. The choice of *ρ* has a strong influence on the resulting estimate and we want to choose as small of a value of *ρ* as the data will allow. Using the representer theorem, which states that a solution to the penalized log likelihood function

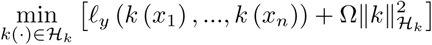

takes the form

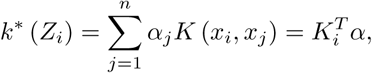

[17]

then the penalized log likelihood function can be rewritten as

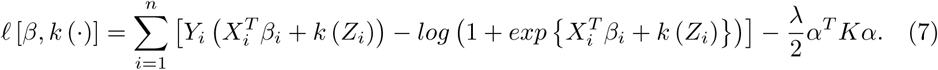

Solving for *α* and *β* gives the closed form equations

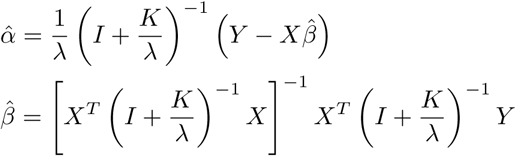

and then, plugging in 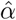 into *k\** (*Z_i_*),

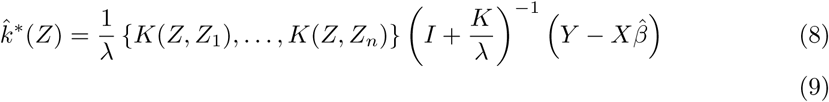

[15].

However, it is possible to approach the penalized likelihood function, *ℓ*[*β, k* (·)], from a generalized linear mixed models (GzLMM) perspective. As logistic regression is a special case of GzLMM, the kernel estimator within the semiparametric logistic regression model parallels the penalized quasi-likelihood function from a logistic mixed model, letting *τ* = 1/*λ* denote the regularization parameter and *ρ* remaining the bandwidth parameter [15]. These parameters can be treated as variance components, where *k* (·) ~ *N* (0, *τK*(*ρ*)) can be treated as a subject-specific random effect and the covariance matrix *K* (*ρ*) is an *n* × *n* kernel matrix as previously defined [16]. This then means that estimating *β* and *k* (·) can be done by maximizing the penalized log likelihood

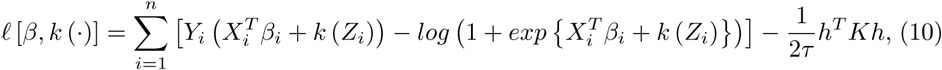

where *h* = *Kα* and *τ* = 1/*λ* [16]. This provides an advantage as it allows for testing of the null hypothesis *H_0_*: *τ* = 1/*λ* = 0 without explicit specification of the basis functions of *f* (·).

However, under the null hypothesis, the kernel matrix *K* disappears, which makes *ρ* a nuisance parameters that is inestimable under the null hypothesis. Davies studied the issue of a nuisance parameter disappearing under the null hypothesis [18], and proposed a score test be used. The score statistic is treated like a nuisance parameter-indexed Gaussian process. To implement this, we must reexamine the likelihoods in Eq 7 and Eq 10. As Eq 7 is a nonlinear function of (*α, β*), Newton-Raphson needs to be implemented to maximize Eq 7 in terms of (*α, β*). If (j) is the *j^th^* iteration of the algorithm, then the (*j* + 1) step solves

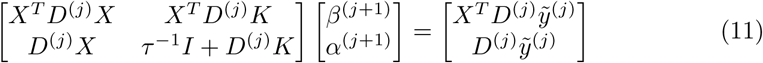

where 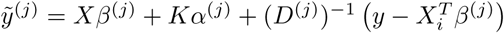, 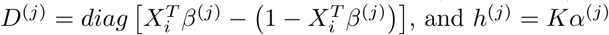, and *h*^(*j*)^ = *Kα*^(*j*)^. Also noting the *β* and *k* (·) depend on *τ* and *ρ,* can be estimated using penalized quasi-likelihood under a logistic mixed model paradigm, then we can rewrite Eq 10 as

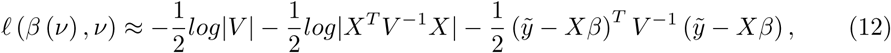

where *v =* (*τ, ρ*) and *V* = *D*^−1^ + *τK.* Then 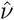 can be solved for in the usual way [16]. However, if the derivative of (9) is taken with respect to *τ*, then the score test for *H*_0_: *τ* = 1/*λ* = 0 can be written as

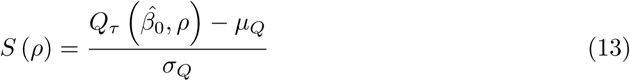

where 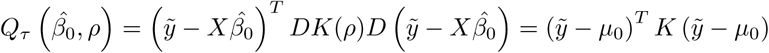, 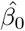 is the maximum likelihood estimate of *β* under the null hypothesis, 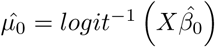, *μ_Q_* = *trace*[*P*_0_*K*(*ρ*)], 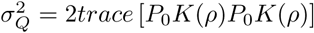, *P*_0_ = *D*_0_ − *D*_0_*X* (*X^T^ D*_0_*X*)^−1^ *X^T^ D*_0_, and *D*_0_ = *diag* [*µ*_0_ − (1 − *µ*_0_)] [16].

*S* (*ρ*) under the null hypothesis is an approximate, *ρ*-indexed Gaussian process, which allows us to apply Davies’ results [18] to get the upper bound for the score test’s p-value. It can be seen that large values of 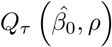 would result in a rejection of *H*_0_, then the upper bound of the p-value is

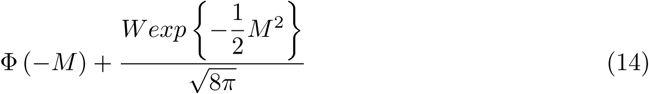

where Φ(·) is the normal cumulative distribution function, *M* is the maximum of *S* (*ρ*) over all of the searched range of *p,* 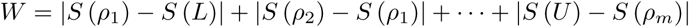, *L* and *U* are the lower and upper bounds, respectively, of the search area for *ρ,* and *ρ_m_* are the search points between *L* and *U* [18] [16]. Liu et al. suggest setting the lower and upper bounds of the *ρ* search to be 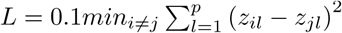 and 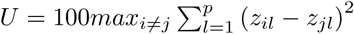 [16].

## Results and Discussion

### Simulations

Functional connectivity matrices were simulated using the `MNS` package in `R`. This package uses the mixed neighborhood selection (MNS) algorithm, which separates a network into two components: edges that are present in the majority of the sample and those that are subject-specific [19]. Using the preferential attachment model proposed by Barabási and Albert in 1999, a set of edges, denoted *E^POP^,* is shared across all subjects in the sample, where edge strengths are uniformly sampled [19]. Inter-subject variability among the edges, denoted 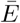, are chosen according to the Erdös and Rényi model, where a choice of the number of elements of 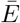 determines the number of random edges and, thus, the level of inter-subject variability [19].

To simulate the data that corresponds to the functional connectivity networks, a multivariate normal distribution is utilized,

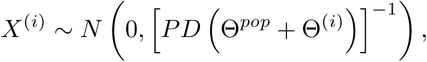

where *PD* (·) is a function that ensures a positive definite standard deviation matrix, Θ^*pop*^ denotes the population networks, and Θ^(*i*)^ denotes the subject-specific networks.

To simulate the functional connectivity data using the `MNS` package, the `gen.Network()` function was called; the parameters associated with the `gen.Network()` are detailed in the `MNS` package documentation [19]. The result is an `S3` object of class `MNS`. Fig 1 below shows an example of simulated functional connectivity networks for three subjects.

**Fig 1.**
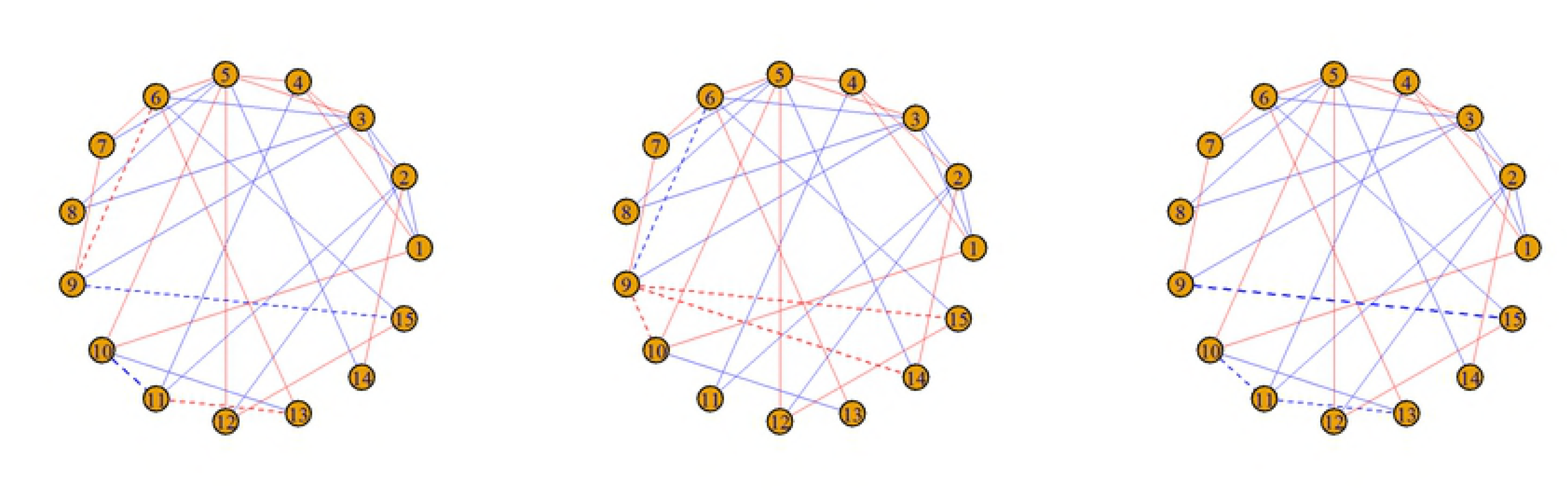
Simulated networks for N=3 subjects under the “cohort” method. Solid lines between nodes represent population edges and dashed lines represent subject-specific edges. Red edges represent a positive association between edges while blue edges represent negative associations.

~~~
> library (MNS)
> set.seed(80045)
> N<-3
> Nets<–gen. Network (method=” cohort” ,p=15,Nsub=N, sparsity=0.2,
REsize=10,REprob=0.5, REnoise=1)
> plot (Nets, view=” sub”)
~~~

For all simulations, the following settings were utilized in the `gen. Network ()` function, which seem to best represent the variability in functional connectivity networks from resting state MRI datasets: `p=90, sparsity=0.75, REsize=10, REprob=0.65, REnoise=3`.

However, the data simulated in the `MNS` package does not exactly match data from a resting state fMRI scan. The existence of negative correlations between brain networks has been a hotly contested debate within the neuroimaging community; the origin, interpretation, and link to the underlying structural connectivity are still unresolved issues. Because of this, the norm within the field is to zero out any negative correlations within the connectivity matrices before further analysis is performed. Other options include taking the absolute value or to normalize the correlations to be between 0 and 1, although these are far less popular. For the purposes of this manuscript, simulated datasets from the MNS package had all negative correlations either zeroed out or normalized between 0 and 1.

The four properties of a distance - edge importance, weight awareness, edge-“submodularity,” and focus awareness - were tested for the resistance perturbation distance under varying simulation models. Under the edge importance property, in weighted graphs, changes that created disconnected components should be penalized more than changes that maintain the connectivity properties of the graph. We simulated this property by breaking up the simulated connectivity matrix into four, equally-sized quadrants and then zeroing out all non-zero cells within the off-diagonal (quadrants I and III) to create two disconnected components. Then, the same number of components were randomly zeroed out to create a comparable “random” graph. Pairwise RPDs were plotted for 1000 iterations in Fig 2 and summary statistics of the RPDs in Table 1, both below.

**Table 1.** Summary statistics of edge importance property simulations.

| Simulation Paradigm | Minimum | 1st Quartile | Median | Mean | 3rd Quartile | Maximum |
| --- | --- | --- | --- | --- | --- | --- |
| RPD for Disconnected Components | 0.00 | 51.28 | 76.34 | 94.38 | 117.50 | 490.50 |
| RPD for Random Deletions | 0.00 | 0.00 | 14.79 | 24.16 | 27.57 | 914.40 |

**Fig 2.**
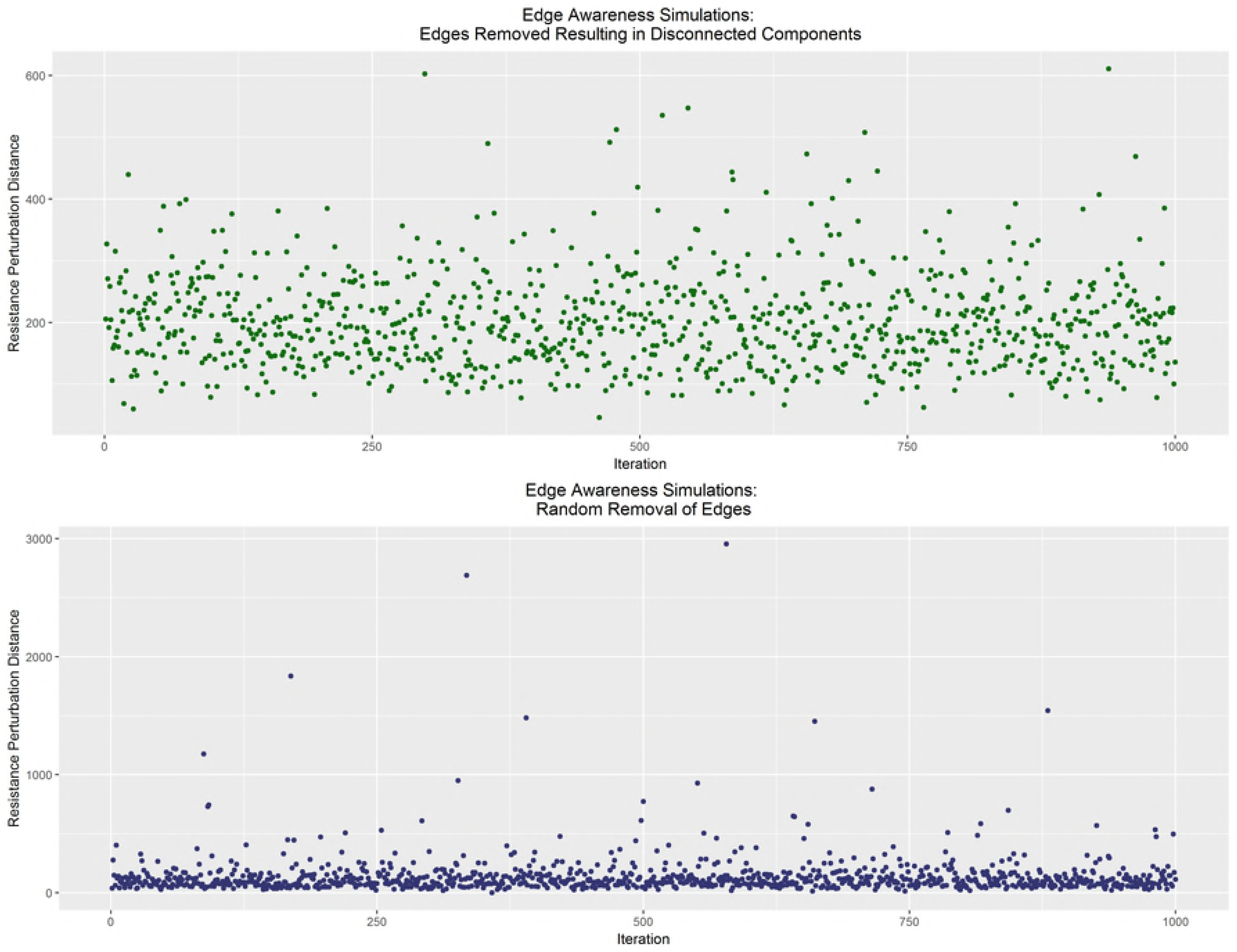
Edge importance property. 1000 iterations under targeted deletions resulting in disconnected components (top) and equally-numbered but random deletions (bottom)

Under the weight awareness property, in weighted graphs, the larger the weight of the removed edge, the greater the impact on the distance. We considered iteratively zeroing out the minimum or maximum non-zero correlation from a simulated ten node connectivity matrix. Pairwise RPDs were plotted for 1000 iterations in Fig 3 and summary statistics of the RPDs in Table 2, both below.

**Table 2.** Summary statistics of weight awareness property simulations.

| Simulation Paradigm | Minimum | 1st Quartile | Median | Mean | 3rd Quartile | Maximum |
| --- | --- | --- | --- | --- | --- | --- |
| RPD for Min Correlation | 2.16 | 20.14 | 43.03 | 77.09 | 103.20 | 583.20 |
| RPD for Max Correlation | 6.44 | 63.64 | 94.26 | 110.90 | 142.80 | 473.60 |

**Fig 3.**
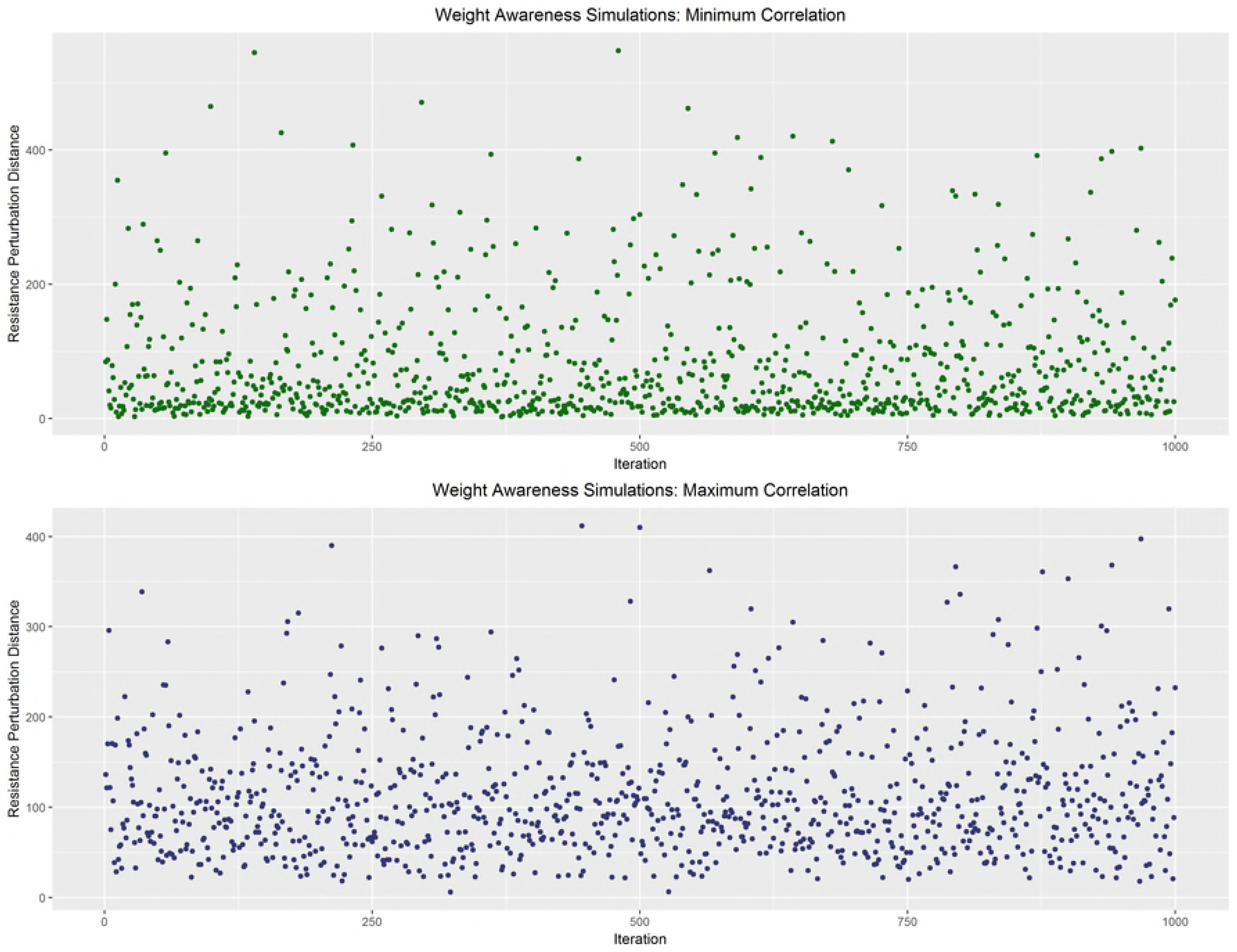
Weight awareness property. 1000 iterations under minimum (top) and maxiumum (bottom) correlation paradigms

Under the edge-“submodularity” property, in weighted graphs, a specific change is more important in a graph with few edges than in a much denser, but equally sized, graph. For ten node graphs, the maximum non-zero correlation was systematically removed from each iteratively-simulated connectivity matrix. Within the `gen.Network()` function, sparsity parameters of 0.45 (for a sparse graph) and 0.95 (for a dense graph). Pairwise RPDs were plotted for 1000 iterations in Fig 4 and summary statistics of the RPDs in Table 3, both below.

**Table 3.** Summary statistics of edge-“submodularity” property simulations.

| Simulation Paradigm | Minimum | 1st Quartile | Median | Mean | 3rd Quartile | Maximum |
| --- | --- | --- | --- | --- | --- | --- |
| RPD for Sparse Graph | 5.31 | 61.53 | 94.73 | 105.00 | 140.60 | 350.50 |
| RPD for Dense Graph | 7.58 | 26.87 | 40.43 | 54.60 | 65.32 | 388.40 |

**Fig 4.**
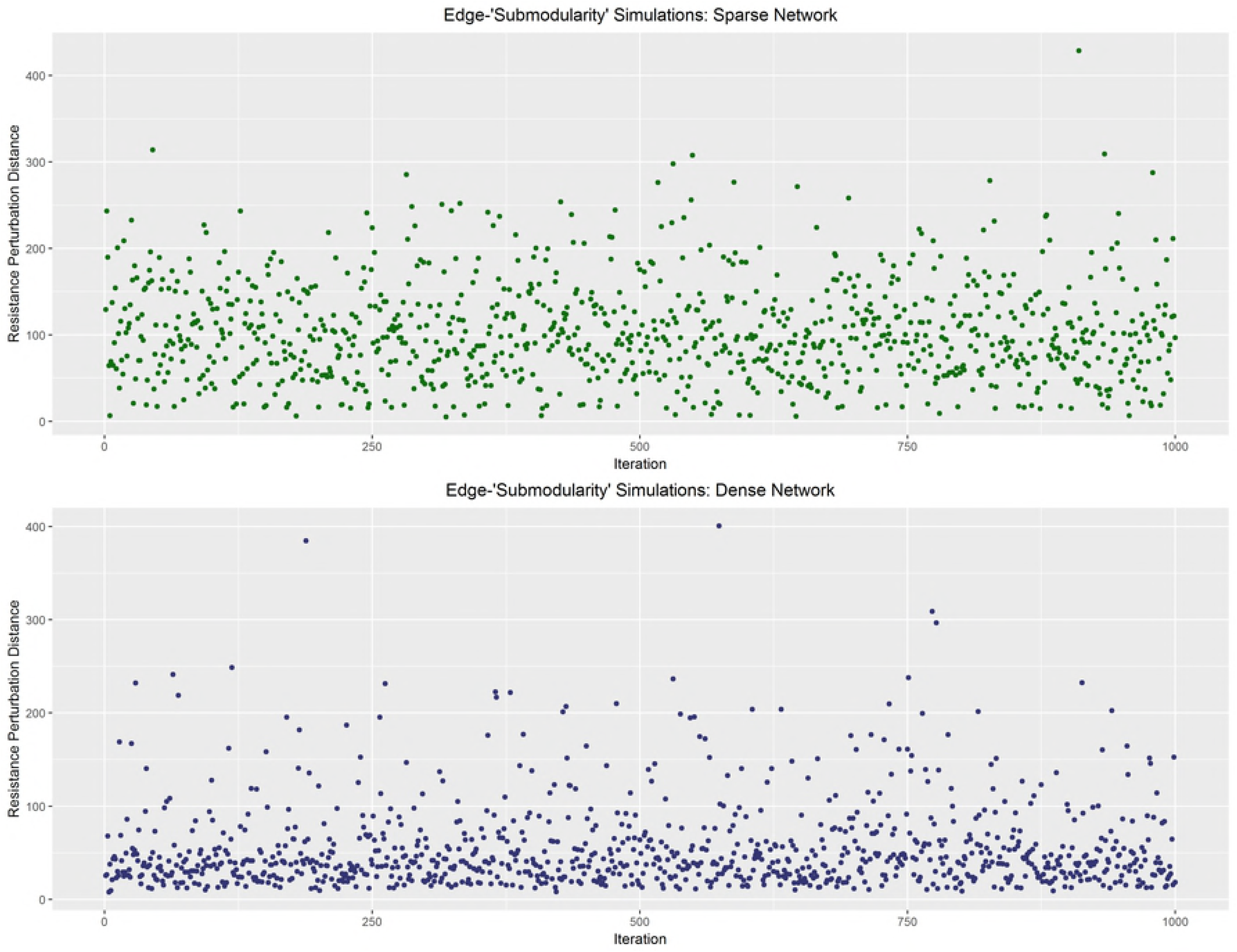
Edge-“submodularity” property. 1000 iterations under sparse (top) and dense (bottom) graph paradigms

Finally, under the focus awareness property, in weighted graphs, random changes in graphs are less important than targeted changes of the same extent. Similar to Koutra et al. [14], for ten node graphs, targeted changes were made by deleting all edges from a randomly chosen node while random changes were made by randomly removing the same number of edges from the whole graph. Pairwise RPDs were plotted for 1000 iterations in Fig 5 and summary statistics of the RPDs in Table 4, both below. As these figures and tables show, the Koutra et al. properties are all satisfied under the simulated constraints of fMRI data.

**Table 4.** Summary statistics of focus awareness property simulations.

| Simulation Paradigm | Minimum | 1st Quartile | Median | Mean | 3rd Quartile | Maximum |
| --- | --- | --- | --- | --- | --- | --- |
| Targeted Changes | 0.00 | 51.08 | 70.45 | 92.95 | 114.40 | 459.50 |
| Random Changes | 0.00 | 0.00 | 14.27 | 24.78 | 29.21 | 456.70 |

**Fig 5.**
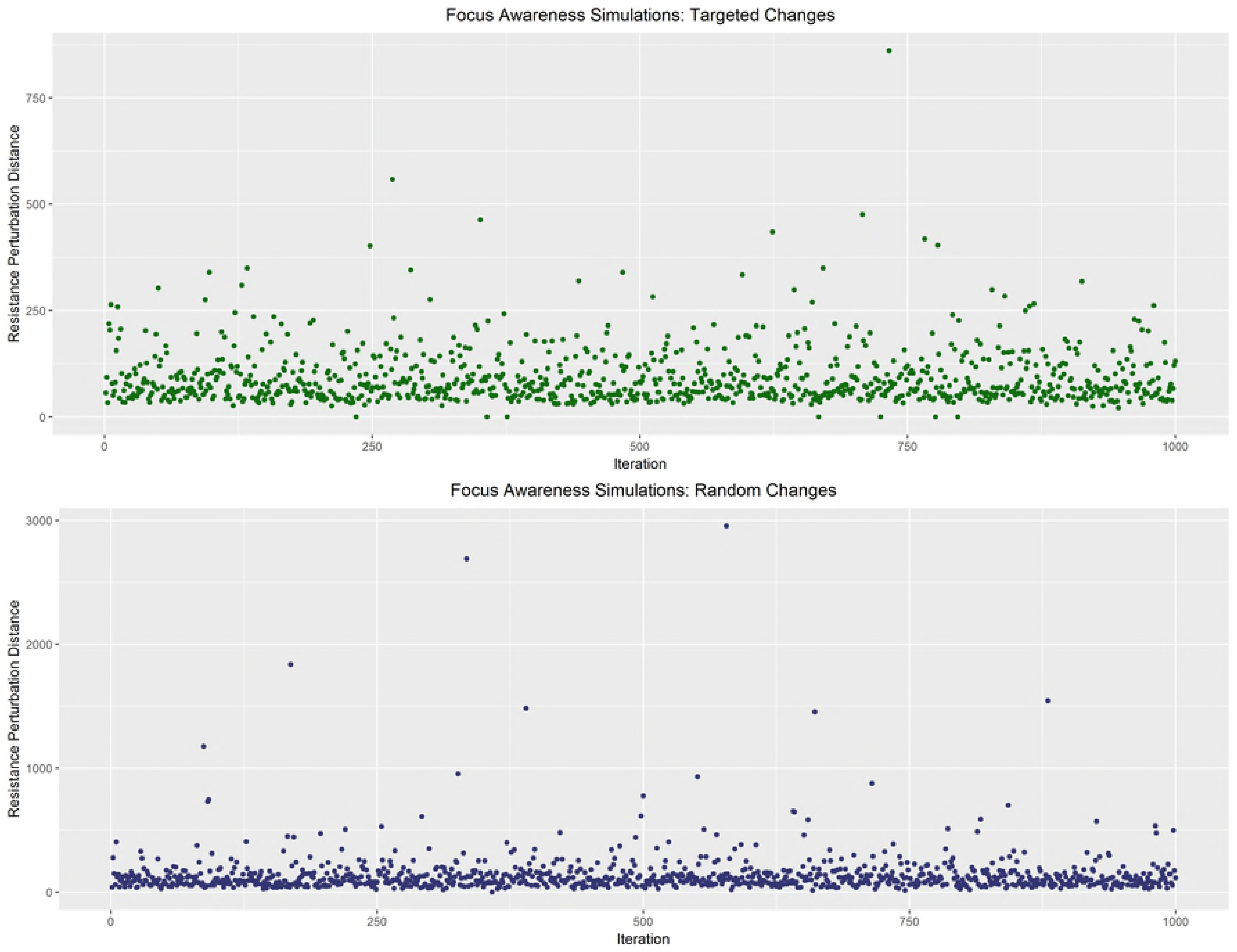
Focus awareness property. 1000 iterations under targeted (top) and random (bottom) change paradigms

Next, to analyze the robustness of the kernel-based score test described, several simulations were conducted. These simulations were split between whether two or one groups of functional connectivity matrices were generated; a simulation under a one generation process presumes that the null hypothesis of all subjects come from the same population is true while a two generation process presumes that the null hypothesis is false and subjects come from two distinct populations. To simplify the analyses for these simulations, no covariates were generated.

We next conducted a series of simulation studies to evaluate the performance of the kernel-based score test under the hypothesis test of *H*_0_: *k* (·) = 0 versus *H_A_*: *k* (·) ≠ 0. As there is no closed-form solution for the test statistic’s accompanying p-value, power and Type I error were calculated using simulated datasets. For the power simulation, 100 different datasets were produced, ten from the “control” population, ten from the “patient” population, and the remaining 80 from a third “noise” population. Each of these populations were simulated under different `gen.Network()` function calls in order to prevent common preferential attachment model parameters. This third population was included to mimic the noisy nature of fMRI data; even among two groups that have been shown to have significant differences in their fMRI data, the vast majority of the graphs are indistinguishable because of the universal way the human brain communicates with itself. The noise population was distributed between the “control” and “patient” populations such that the final sample sizes were 55 in the “control” population and 45 in the “patient” population. Each simulated connectivity matrix was generated with the following parameters: `p=90, sparsity=0.75, REsize=10, REprob=0.65, and REnoise=3`. Bounds of the *ρ* search were set based on the suggestion from Liu et al. [16]. An indicator function was used to determine whether each simulation’s resulting p-value was greater than *α* = 0.05, then the ultimate power of the kernel score test was calculated using 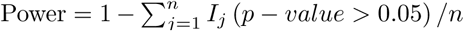, where *n* is the number of repetitions. After 1000 iterations, the empirical power of the kernel-based score test was 0.945. Similarly, for the test statistic’s Type I error rate, all 100 simulated samples came from a single generation process with the same parameters as the power simulation and bounds of the *ρ* search and the final Type I error was calculated using 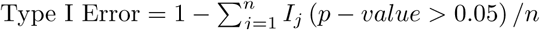, where *n* is the number of repetitions. After 1000 iterations, the empirical Type I error rate of the kernel-based score test was 0.0496. Our simulations show that the empirically-calculated Type I error is very close to the nominal value of 0.05 while the power of our score test has high power to detect true differences in a dataset.

We also conducted a simulation under a two generation process to understand the impact of how the allocation of the “noise’” population to the “control” and “patient” populations affected the kernel-based score test’s p-values. In a very similar manner to the power simulation, three datasets were produced. However, the allocation of the simulated noise population to the control and patient populations was varied; the percentage of the noise population allocated to the control population varied along (5%,95%) by increments of 5%. One hundred iterations occurred at each noise allocation. Any iteration that resulted in a p-value greater than 0.05 was considered a false negative as the simulation was set up in such a way that the underlying truth was that the “control” and “patient” populations were different from one another. The results of the simulation are plotted in Fig 6, below. The highest Type II errors occurred between noise splits where 40% and 55% of the noise was allocated to the control population; this is not surprising as a noise allocation that was nearly evenly split between the two groups would make their average graph look exceedingly similar.

**Fig 6.**
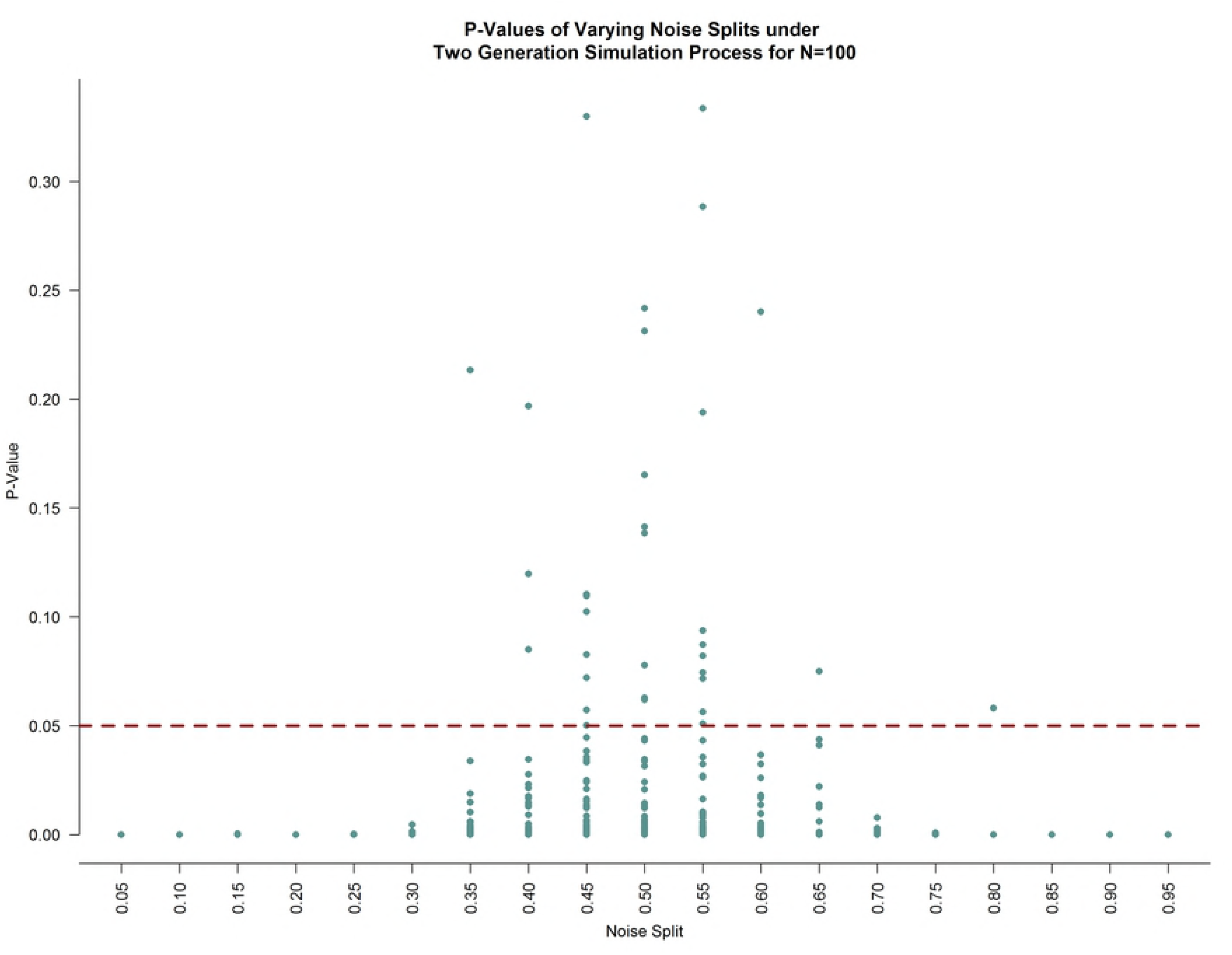
P-values under varying noise population allocations under a two generation simulation process. At each noise allocation, 100 iterations were conducted. The red dashed line is at the nominal p-value of 0.05. Any point above this red line is a false negative.

Similarly, we conducted a simulation under a one generation process to understand how splitting the 100 connectivity matrices between “control” and “patient” populations affected the p-values of the kernel-based score test. Like the Type I error simulation, only one dataset was produced. However, how this dataset was split between the two populations was varied; the number of connectivity matrices allocated to the “control” population varied from five to 95, increasing in increments of one at each iteration. Any iteration that resulted in a p-value less than 0.05 was considered a false positive as the simulation was set up in a way that the underlying truth was that there was no significant difference between “controls” and “patients.” The results of the simulation are plotted in Fig 7, below. The total Type I error across all allocations was 0.05, on par with the empirical Type I error calculated.

**Fig 7.**
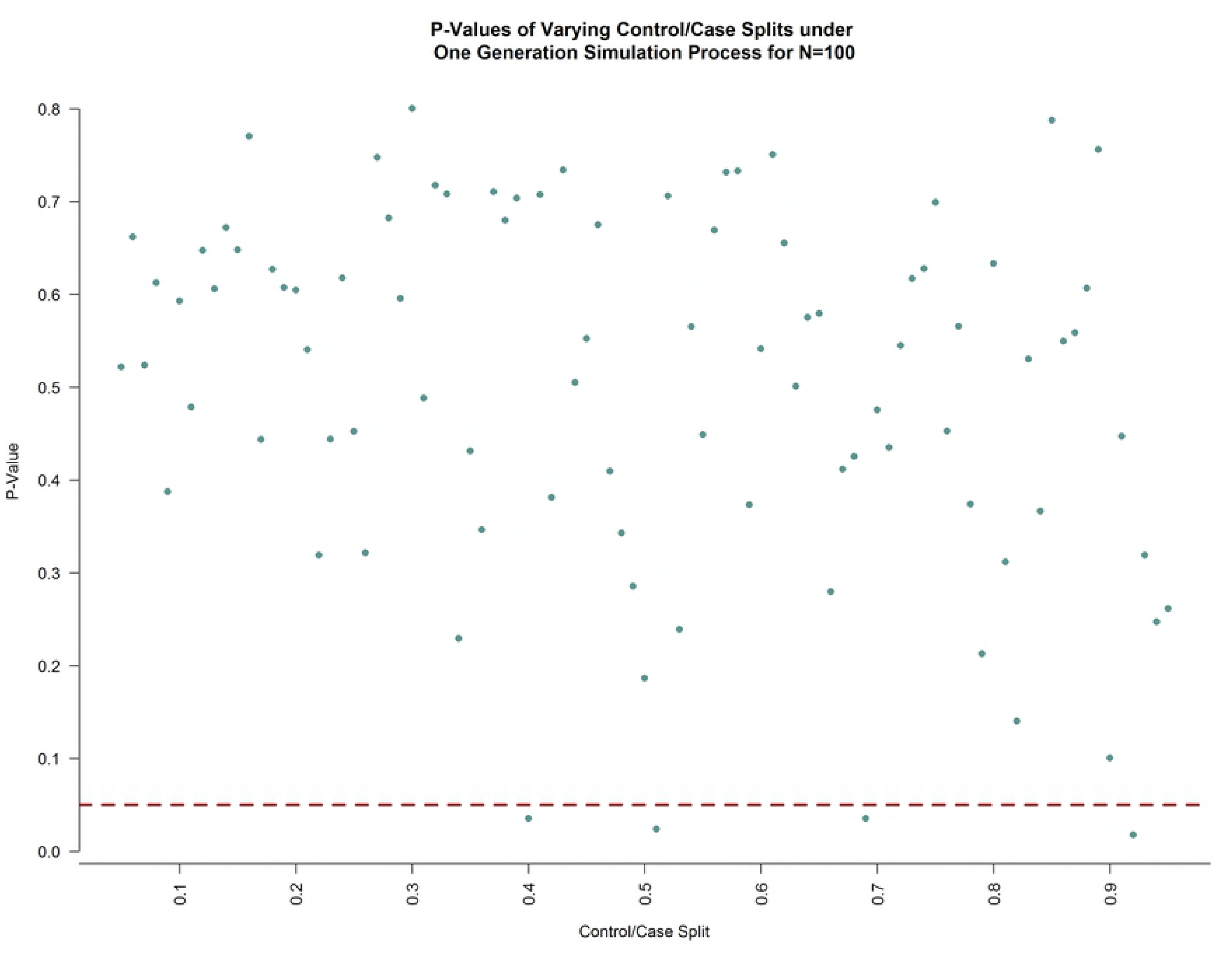
P-values under varying noise population allocations under a one generation simulation process. Allocation of the 100 generated connectivity matrices varied from 5:95 to 95:5, increasing in increments of one. The red dashed line is at the nominal p-value of 0.05. Any point below this red line is a false positive.

Finally, the lower and upper bounds of the grid search for the *ρ* parameter of the kernel were varied under both one- and two-generation processes. While Liu et al. [16] provides a justification for these bounds, because our kernel-based score test is more complex (i.e., it involves semi-parametric estimation of phenotypic parameters and utilizes a different kernel), we wanted to test the robustness of the test under differing *ρ* grid searches. A lower value of 0.00001 and upper value of 0.01, each iterative multiplied by 10*^i^* for *i* = 0, *…*, 10 were multiplied by 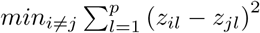 and 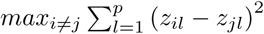, respectively. A total of 100 simulations were conducted under each of the one- and two-generation processes and p-values were plotted. Any p-value that was less than 0.05 under the one-generation process was considered a Type I error while any p-value over 0.05 in the two-generation process was considered a Type II error. Fig 8, below, shows a scatterplot of these simulations. These simulations show that, under a one-generation process, the false positive rate varies according to the boundaries chosen but, overall, is slightly above the empirical Type I error rate at 0.06. Interestingly, for a two-generation process, the boundaries of the *ρ* grid search are highly robust, with the Type II error at 0.00.

**Fig 8.**
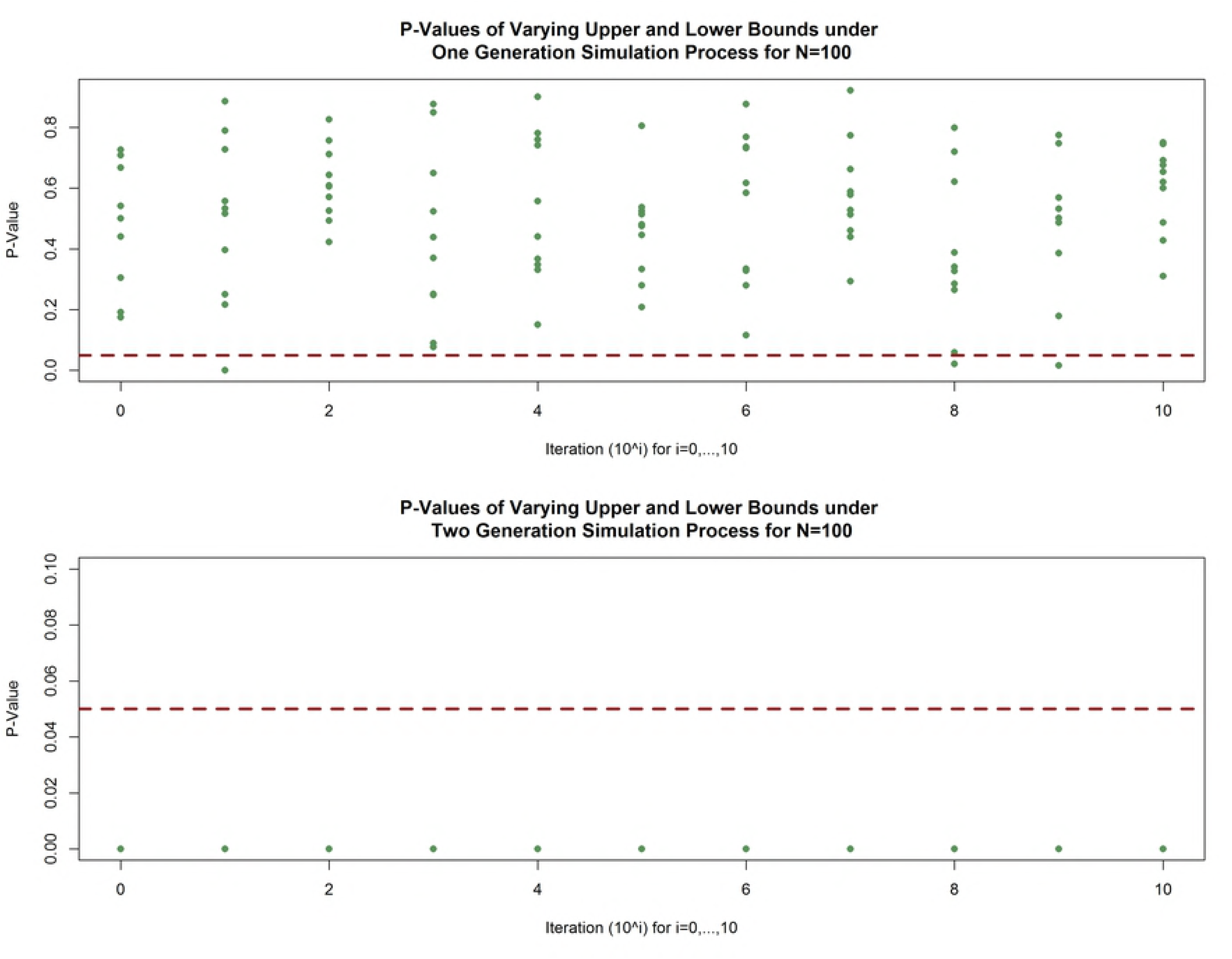
P-values under varying boundaries of the *ρ* parameter. For each of the one generation (top) and two generation (bottom) processes, the minimum and maximum sums of squares from the RPD matrix were multiplied by 0.00001 × 10*^i^* and 0.01 × 10*^i^* for *i* = 0,1, *…*, 10, respectively. Ten iterations at each value of *i* were calculated. Under the one-generation process, any p-value less than the red dashed line - the nominal p-value of 0.05 - is a false positive. Under the two generation process, any p-value greater than the red dashed line is a false negative.

## COBRE Dataset

The Center for Biomedical Research Excellence through the Mind Research Network for Neurodiagnostic Discovery (MRN) is one of many National Institute of Health (NIH)-funded institutions that focus on the development of “new, disease-specific research centers or augment[ing] the capability of existing centers [20].” The MRN’s focus is on the neural mechanisms of schizophrenia, with the unifying theme being that the condition is characterized by “structural, functional, and effective connectivity between cortical and subcortical brain regions, producing abnormalities in the integration of information across distributed brain circuits [21].” Previous studies [22] [23] of this dataset have shown significant differences between schizophrenia and control patients in the hippocampus and default mode network (a large scale brain network comprised of medial prefrontal cortex, posterior cingulate cortex, and inferior parietal lobule) with more subtle differences in the temporal and frontal networks. However, neither of these studies approached their analysis from a graph theoretic perspective, choosing instead to perform versions of a mass univariate analysis.

As its contribution to the 1000 Functional Connectomes Project, the MRN contributed raw anatomical and functional MR data from 72 patients with diagnosed schizophrenia and 75 healthy controls, although two control patients had to be excluded due to disenrollment [24]. A multi-echo, magnetization prepared rapid gradient echo (MPRAGE) sequence was used to acquire the anatomical information on each subject, with the following parameters: TR/TE/TI=2530/[1.64, 3.5, 5.36, 7.22, 9.08]/900ms; flip angle=7°; FOV=256×256mm; slab thickness=176mm; matrix=256×256×176; voxel size=1×1×1mm; number of echoes=5; pixel bandwidth=650Hz, total scan time=6 minutes [24]. Resting state functional MR data was acquired using a single-shot, full k-space, echo planar imaging (EPI) with ramp sampling correction using the intercommissural line as the reference and the following parameters: TR=2 seconds; TE=29ms; matrix size=64×64; 32 slices; voxel size=3×3×4mm [24]. In addition to this imaging data, the MRN also provided phenotypic information on each subject, including age, gender, handedness, and diagnostic information, when applicable. Table 5 provides a summary of the phenotypic information on controls and patients.

**Table 5.** COBRE Dataset Subject Demographics.

|  | Control (n=73) | Patient (n=72) |
| --- | --- | --- |
| Age in years, mean (SD) | 35.90 (11.64) | 38.17 (13.89) |
| Sex, n (%) |  |  |
| Male | 50 (68.5%) | 58 (80.56%) |
| Female | 23 (31.5%) | 14 (19.44%) |
| Handedness, n(%) |  |  |
| Left | 1 (1.35%) | 10 (13.89%) |
| Right | 71 (95.94%) | 60 (83.33%) |
| Ambidextrous | 2 (2.70%) | 2 (2.78%) |

An automated pre-processing and denoising pipeline was implemented with the CONN software package within MatLab [25]. Within this pipeline, the first four volumes were discarded to ensure T1 equilibrium effects, each subject’s images were realigned to the first volume, but no slice-timing correction was applied as images were acquired in a descending manner. Data were spatially normalized to the Montreal Neurological Institute (MNI) space and smoothed using a Gaussian kernel with a full-width at half-maximum of 8mm. During the denoising process, two different sources of possible confounds were regressed out: (1) BOLD signal from white matter and cerebrospinal fluid (CSF); and (2) realignment parameters (6 parameters). Correlation matrices were then extracted from CONN following a first-level ROI-to-ROI analysis. A hybrid physical atlas was used, where the FSL Harvard-Oxford atlas was used to parcellate the cortical and subcortical areas and the Automated Anatomical Labeling (AAL) atlas [26] was used to to parcellate the cerebellar areas; this resulted in a physical atlas of 132 regions. Weighted networks were extracted from the `*.mat` files using the `R.matlab` package. As the matrices contained Fisher’s transformed correlation coefficients, the hyperbolic tangent function was applied to all correlations then negative correlations were set to zero.

Using the entire dataset, which included 72 schizophrenia and and 73 control patients following the pre-processing and denoising procedures, the outcome was a binary classification variable of schizophrenia diagnosis. The regression parameters for the phenotypic covariates of age, sex, and handedness were parametrically estimated while the RPD matrix was non-parametrically estimated. Specifically, we considered the following semiparametric logistic model:

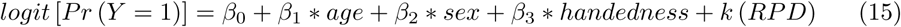

where *k* (·) is a nonparametric kernel distance function of the 132 × 132 RPD matrix. Details of the estimation procedure can be found in the Methods section. Additionally, we also considered a simpler, fully non-parametric logistic model, which does not include any of the phenotypic covariates:

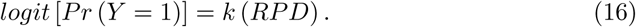

This was done to test whether the phenotypic covariates were confounders in the association between the RPD matrix and binary schizophrenia classification.

The same two models (semiparametric and fully non-parametric) were fit to the full dataset, but for which all negative correlations within the subject-level fMRI connectivity matrices left as is. This was done to determine whether there was a significant loss of information by following the neuroimaging standard of zeroing out any negative correlation between regions of interest. Recent articles [27] [28] have pointed to a potentially significant physiological role of negative correlations within fMRI. Specifically, Parente et al. notes that, while these negatively correlated brain networks still lack a well-defined biological explanation, they appear to be have an association with the alterations in brain function in people diagnosed with schizophrenia [27].

The results of these analyses are present in Table 6, below.

**Table 6.** Analysis of full COBRE dataset.

| Regression | | P-value of score test for $H_0 : k(\cdot) = 0$ |
| --- | --- | --- |
| Semiparametric | Zeroing out Negative Correlations | $p = 0.53$ |
| Semiparametric | Keeping Negative Correlations | $p = 0.61$ |
| Fully Non-Parametric | Zeroing out Negative Correlations | $p = 0.02$ |
| Fully Non-Parametric | Keeping Negative Correlations | $p = 0.48$ |

Table 6 shows that only under the fully non-parametric paradigm when the negative correlations were zeroed out did we reject the null hypothesis of *H*_0_: *k* (*·*) = 0. Our hypothesis that keeping all negative correlations within the dataset would preserve more information was not confirmed as both the semiparametric and fully non-parametric models were not significant at the *α* = 0.05 level. Heat maps of the average connectivity matrix for control versus schizophrenia patients under both paradigms can be seen in Fig 9, below. These heat maps show exceedingly similar patterns of average connectivity between the two groups, which may be the reason why the RPD-based score test was unable to find a significant difference in most of the scenarios we considered.

**Fig 9.**
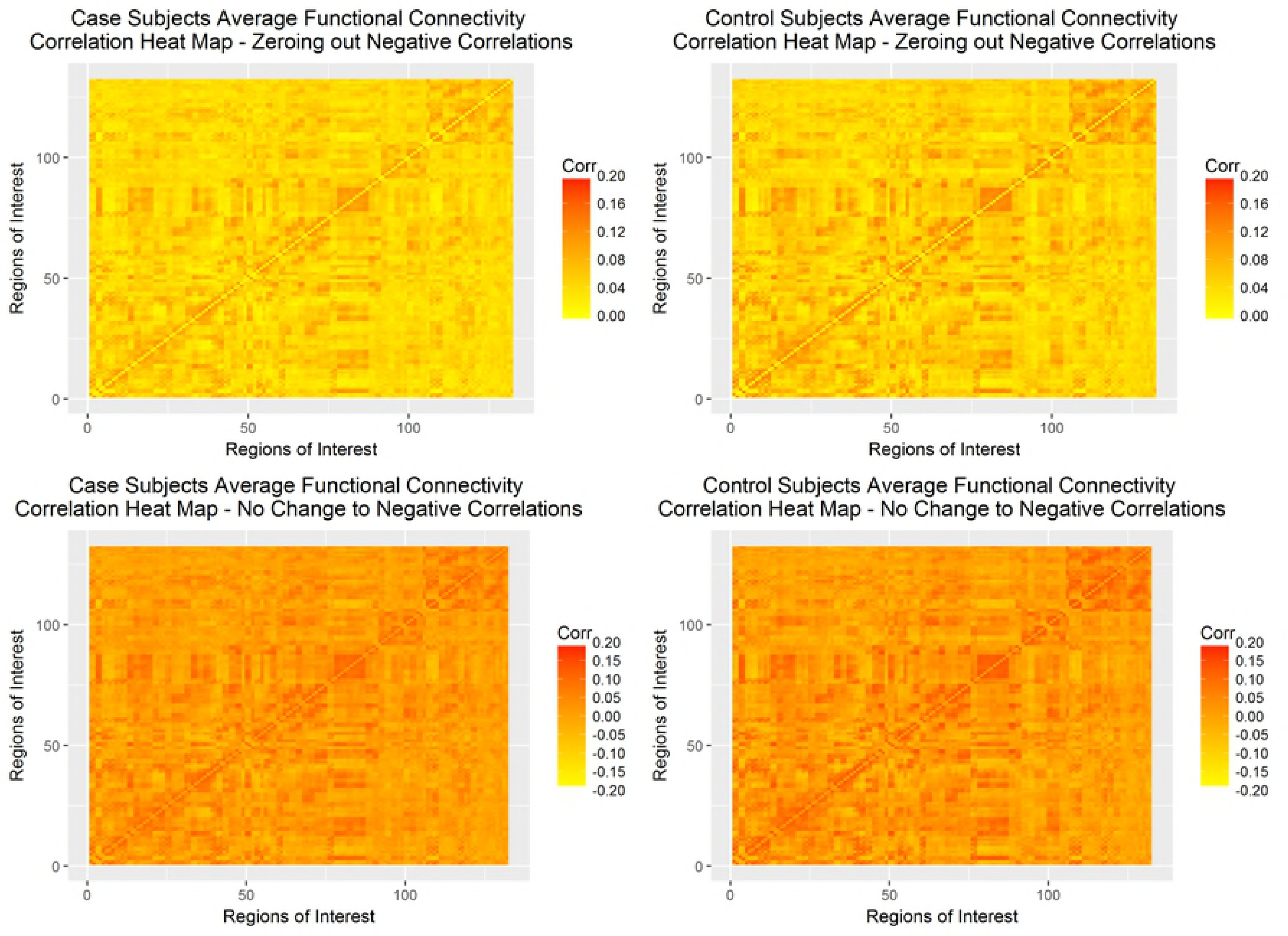
Correlation heat maps full COBRE-I dataset. Correlation heat maps of control and schizophrenia subjects under zeroing out negative correlations (top row) and normalizing all correlations (bottom row).

A subset of the entire COBRE dataset, which included all schizophrenia subjects who had an diagnosis of paranoid schizophrenia (ICD-9 code of 295.3) and an equal number of randomly selected control subjects, was analyzed. To ensure comparable groups, frequency matching for handedness, sex, and age category (18 – 25, 26 – 35, 36 – 45, 46+) was conducted. Because schizophrenia is such a heterogeneous condition, we believed that by restricting our sample of cases to only those with the same sub-diagnosis, we would be removing some of the noise present within the dataset exogenous to the normal variation in fMRI connectivity. As with the full dataset, four different regression models were fit to the data, the results of which are summarized in Table 7, below.

**Table 7.** Analysis of COBRE dataset - Paranoid Schizophrenia Cases Only.

| Regression | | P-value of score test for $H_0 : k(\cdot) = 0$ |
| --- | --- | --- |
| Semiparametric | Zeroing out Negative Correlations | $p = 0.49$ |
| Semiparametric | Keeping Negative Correlations | $p = 0.59$ |
| Fully Non-Parametric | Zeroing out Negative Correlations | $p = 0.15$ |
| Fully Non-Parametric | Keeping Negative Correlations | $p = 0.46$ |

Table 7 shows that for all four conditions, we fail to reject the null hypothesis of *H*_0_: *k* (*·*) = 0. Heat maps of the average connectivity matrix for the randomly-selected control versus paranoid schizophrenia patients under both paradigms can be seen in Fig 10, below. As with the full COBRE dataset, the heat maps do not show differences in the totality of functional connectivity. However, in both cases, the schizophrenia patients appear to have lower correlations between regions of interest than the control patients.

**Fig 10.**
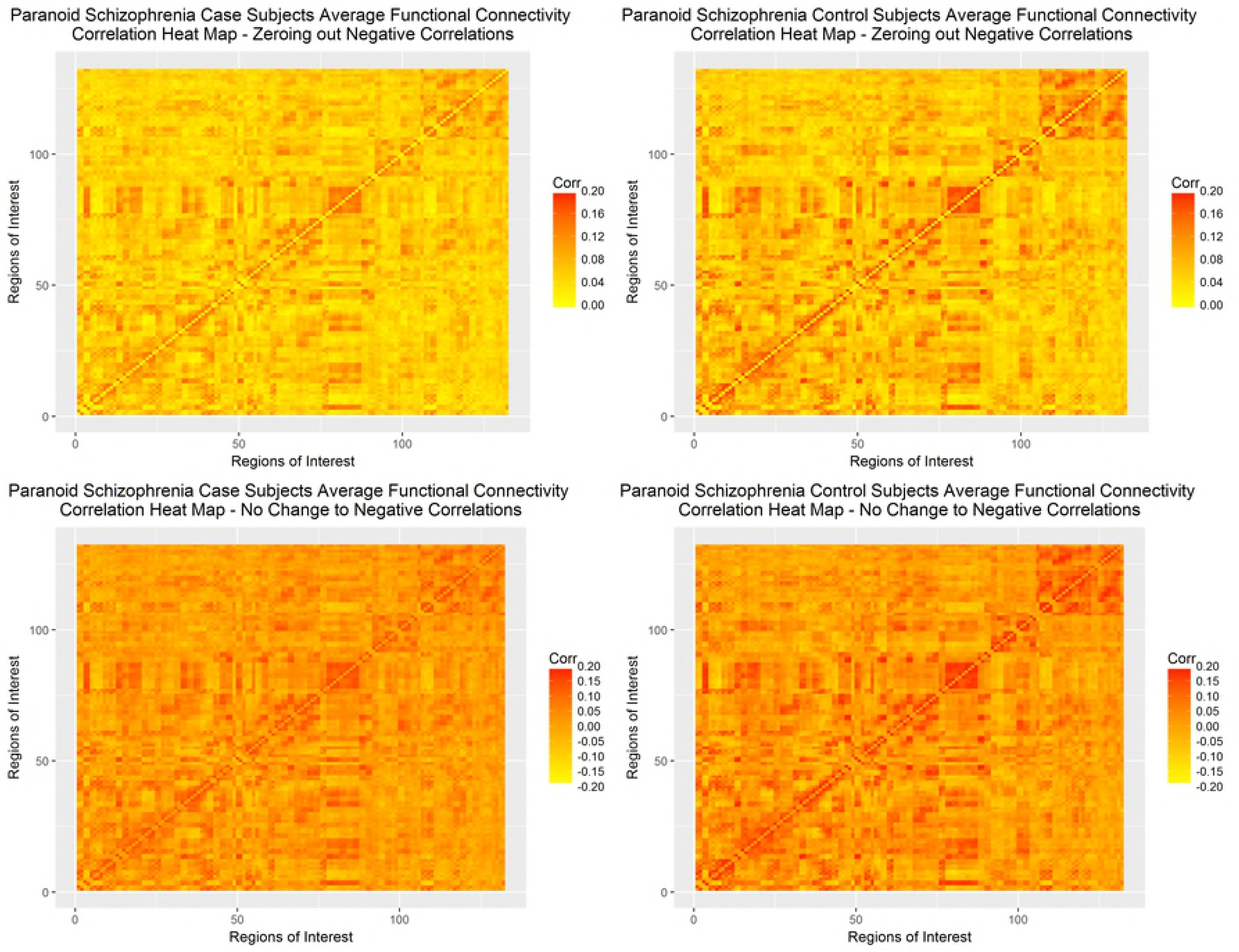
Correlation heat maps paranoid schizophrenia COBRE-I dataset. Correlation heat maps of control and schizophrenia subjects for only the paranoid schizophrenia subset under zeroing out negative correlations (top row) and normalizing all correlations (bottom row).

## Conclusion

In this paper, we applied a concept from the electrical engineering field, the resistance perturbation distance, to a kernel logistic regression framework, where the outcome of interest is a binary classifier, phenotypic covariates are modeled parametrically, and the distance metric is modeled nonparametrically using a kernel machine method. The RPD is computationally efficient and does not result in a loss of information on either a local or global scale, unlike many other graph theoretic measures. The application of a kernel logistic regression allows for the RPD to be modeled without making any assumption as to the parametric form of its association with the binary classifier. Because our model is semi-parametric, we are able to control for potential phenotypic confounders within a parametric framework, allowing for ease of parameter estimate interpretation, should they be desired. Further, the kernel regression framework could be extended to account for repeated measures, allowing for RPD metrics to be calculated at multiple points during each subject’s fMRI scan time.

There are several limitations that affect our approach. First, while our model proved to have high power, a low Type I error rate, and is robust to varying study design and searchable spaces of the score test statistics, only one significant association was found between the RPD matrix and the binary classifier in the full COBRE-I dataset under the fully non-parametric score test with negative correlations zeroed out. However, when accounting for multiple comparisons, this association is no longer significant. The difference between simulation and real datasets could be due to a variety of factors, either working in isolation or compounded on one another. Several recently-published studies [8] [29] [30] have noted that choice of pre-processing pipeline can impact the results of an inferential analysis involving graph theoretic measures, especially in resting state fMRI. We have not studied the impact of different parameters within the same pre-processing pipeline nor the impact of an entirely different manner of pre-processing on the RPD. As well, it was noted earlier that, while the overall patterns of connectivity within the heat maps appear to be similar between cases and controls, the overall magnitude of the correlations may differ; the RPD is not sensitive to a global difference in the magnitude of edge weights as it is scale invariant. Finally, while no self loops allows for desirable mathematical properties of simple graphs, its absence is significant biologically. Network function is maintained by biologic feedback loops, which cannot be modeled with the current graph theory framework. These feedback loops could have particular importance in the distinction of fMRI connectivity patterns between controls and those with schizophrenia.

A future direction within this modeling approach could be to use the RPD and kernel logistic regression within a more confined brain atlas. The hybrid atlas contains 132 parcellated regions covering the entirely of the brain. However, it may be that restricting this methodology to pre-specified regions of interest may bear results more comparable to that seen in simulation. Additionally, as the RPD is scale invariant, relative, rather than absolute, differences in connectivity may be more informative for this algorithm. Specifically, if the total sample average connectivity between two nodes is some value *ρ* satisfying −1 *< ρ <* 1 then looking at differences in individual deviation values from this average, rather than the absolute differences, may help circumvent the scale invariance of the distance metric. Finally, an extension to take into account repeated measures could help provide an analytical framework for analyzing resting state fMRI as rs-fMRI connectivity is known to not be consistent across the entirely of the scan time.

## Acknowledgments

We thank Michael Regner, M.D. for his insight into the use and troubleshooting of the CONN toolbox within MatLab as well as providing comments that greatly improved the manuscript.

